# Infection triggers tumor regression through activation of innate immunity in *Drosophila*

**DOI:** 10.1101/552869

**Authors:** Camille Jacqueline, Jean-Philippe Parvy, Dominique Faugère, François Renaud, Dorothée Missé, Frédéric Thomas, Benjamin Roche

## Abstract

The pioneering work of Dr. William Coley has shown that infections can stimulate the immune system and improve tumor growth control. However, the immune mechanisms responsible for the protective role of infectious agents have still not been identified. Here, we investigated the role of innate immune pathways in tumor regression by performing experimental infections in genetically modified *Drosophila* that develop invasive neoplastic tumors. After quantifying tumor size, through image processing, and immune gene expression with transcriptomic analyses, we analyzed the link between tumor size and pathogen-induced immune responses thanks to a combination of statistical and mathematical modeling. *Drosophila* larvae infected with a naturally-occurring bacterium showed a smaller tumor size compared to controls and fungus-infected larvae, thanks to an increase expression of Imd and Toll pathways. Our mathematical model reinforces this idea by showing that repeated acute infection could results in an even higher decrease in tumor size. Thus, our study suggests that infectious agents can induce tumor regression through the alteration of innate immune responses. This phenomenon, currently neglected in oncology, could have major implications for the elaboration of new preventive and immunotherapeutic strategies.

**One Sentence Summary:** Bacterial infections can decrease cancer cell accumulation through stimulation of innate immune responses.

## Introduction

Most, if not all, multicellular organisms are exposed, during their life, to a diversity of infectious agents (through contact, ingestion and inhalation, among other possibilities) that stimulate the innate immune system (*1*) but also let an imprint on individual’s immune profile (*2*). As numerous immune mechanisms are shared by anti-infection and anti-cancer responses (*3*), it is theoretically expected that the personal history of infections could influence cancer cell elimination/proliferation in the body. For instance, evidences are accumulating to suggest that infectious agents may play an indirect role in tumor progression, through immunological, ecological and evolutionary interactions with immune system (*4*). Moreover, early work from William Coley as well as many other reports have unambiguously linked infection to tumor regression in cancer patients (*5, 6*) suggesting that non-specific immunity may be particularly relevant for cancer treatment (*7*). While the impact of infectious agents on the adaptive tumoral immunity has been studied in mammals (*8, 9*), the activation of innate immune pathways, a non-specific first line of defense, have however been poorly investigated in term of cancer prevention.

To disentangle the role of infection-induced innate immunity in tumor burden, we performed our experiments in an invertebrate model which does not evolve an adaptive immunity (but see (*10, 11*)). In this context, *Drosophila* is recognized as a pertinent model for the study of cancer (*12*). We considered two types of acute infections, bacterial and fungal, since they trigger different immune pathways in an experimental system of *Drosophila*, which develops GFP-labeled invasive neoplastic tumors in larval epithelial tissues called eye-antennal discs (*13*). We combined extensive image analysis with transcriptomic study of the three main immune pathways of drosophila immunity. Then, we used statistical analysis to figure out how immune expression following each kind of infection modulates accumulation of cancer cells. Finally, we developed a mathematical model calibrated with our experimental data to show that the repetitive activation of innate immunity by acute infections may represent a significant factor for cancer prevention.

## Results and Discussion

We first measured the activation of innate immune pathways in response to cancer cells without infection. In *Drosophila,* three main pathways are responsible for driving expression of downstream effectors of innate immunity (*14, 15*). Immune Deficiency (Imd) is involved in the response to Gram-negative bacterial infection (*16*) while Toll pathway is stimulated in response to fungi and gram-positive bacteria (*17, 18*) and Janus Kinase/Signal Transducer and Activator of Transcription (Jak/STAT) is activated by injury or by the presence of aberrant cells (*19, 20*) and upon parasitoid wasps infection (*21*). qRT-PCR analysis revealed no effect of cancerous status on *diptericin* (*dpt*, a read-out for Imd pathway) neither on *unpaired 3* (*upd3*, a key activator of Jak/STAT pathway) expression. However, the presence of tumor cells influenced *drosomycin* expression (*drs*, a read-out for Toll pathway) with cancerous larvae expressing higher level of *drs* than non-cancerous ones (Figure 1A; z-value = 2.04, pval = 0.04). As previously reported, our results suggested that Toll pathway is activated in presence of cancer cells (*22*).

**Figure 1:**
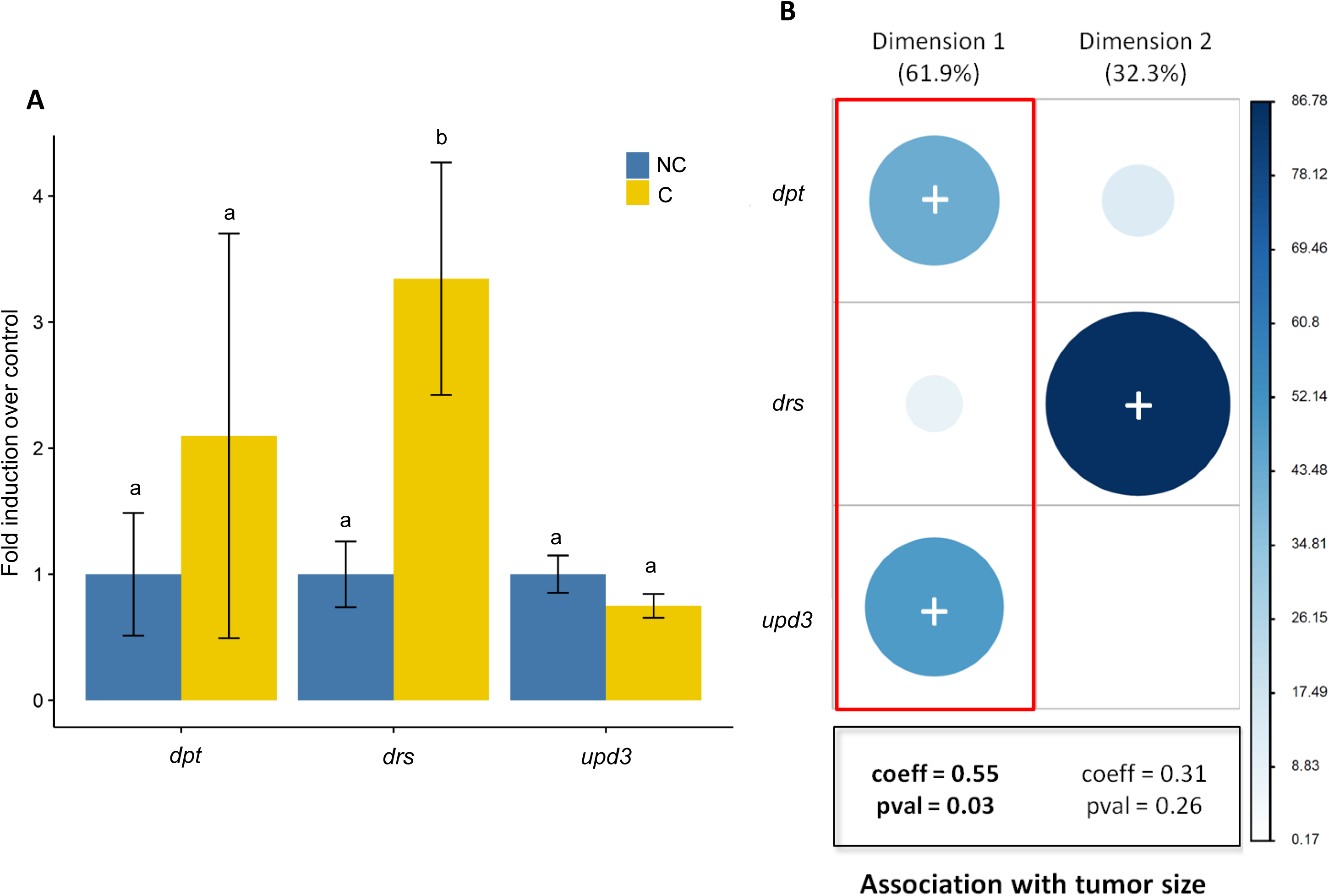
A) Fold induction of the three immune genes in cancerous larvae in yellow without infection relatively to non-cancerous larvae in dark blue. The error bars represent the standard error of mean relative expression calculated for all pools. Shared letters (a, b) above error bars denote means that are not significantly different, while different letters indicate that p<0.05 for pairwise comparisons (Tukey’s test). B) Contribution of the three immune genes in PCA dimensions. The scale shows the intensity of the contribution. Signs in the circle represent the sign of the association. The scare on the bottom reports the results of statistical analyses of model explaining tumor size with the dimensions of the PCA. (Abbreviation : *dpt* : *diptericin*; *drs* : *drosomycin: upd3 : unpaired 3*)

Still with non-infected *Drosophila*, we quantified the tumor size using the number of pixels with a standardized fluorescent intensity above a given threshold (see Supplementary materials for sensitivity on this measure). Using a generalized linear mixed model, we observed that tumor size was associated with the interaction of the three immune pathways (triple interaction: χ^2^ = 4.6, df=1, pval = 0.03). Therefore, we used a principal component analysis (PCA) based on the expression of the three immune genes to study their complex impact on tumor growth. We found that *dpt* and *upd3* were positively and significantly associated to the first dimension, which accounted for 61.9% of the total variance (Figure 1B). *drs* was positively associated to the second dimension, which accounted 32.3% of the total variance (Figure 1B). Statistical analyses showed that tumor size was significantly associated with this first dimension (Figure 1D; t=2.3, df = 13, pval = 0.03). Taken together, these results suggest that the activation of Toll pathway is not proportional to the number of cancer cells but that Imd and Jak/STAT activations are linked with tumor size in non-infected larvae. Accordingly, evidences have shown that *upd3*-dependent Jak/STAT activation in *Ras*^*v12*^*/scrib*^*1*^ tumors promotes tumor growth (*23*). Moreover, activation of Imd pathway has also recently been reported in tumor bearing larvae, mutant for *dlg* or *mxc* (*24, 25*). As we measured gene expression in whole body, and not specifically in tumor tissues, the upregulation of *upd3* may have not been sufficient to be detected in cancerous larvae in our first qRT-PCR analysis.

In order to evaluate the effect of infections on tumor burden, we carried out infectious treatments on tumor bearing larvae using the gram-negative bacterium *Pectobacterium carotovorum carotovorum* (*Pcc*) or the fungal entomopathogen *Beauvaria bassiana* (*Bb*) and compared tumor growth in those conditions with non-infected larvae (see supplement materials Table S1 for prevalence data). Analysis of tumor phenotype showed that larvae orally infected by the bacterium *Pcc* showed smaller sizes compared to non-infected ones (Figure 2A; z-value = 3.5, pval = 0.001). However, infection by *Bb* was not significantly affecting tumor size compared to controls. This suggested that infections may have different effect on tumor regression depending on the characteristics of the downstream immune response.

**Figure 2:**
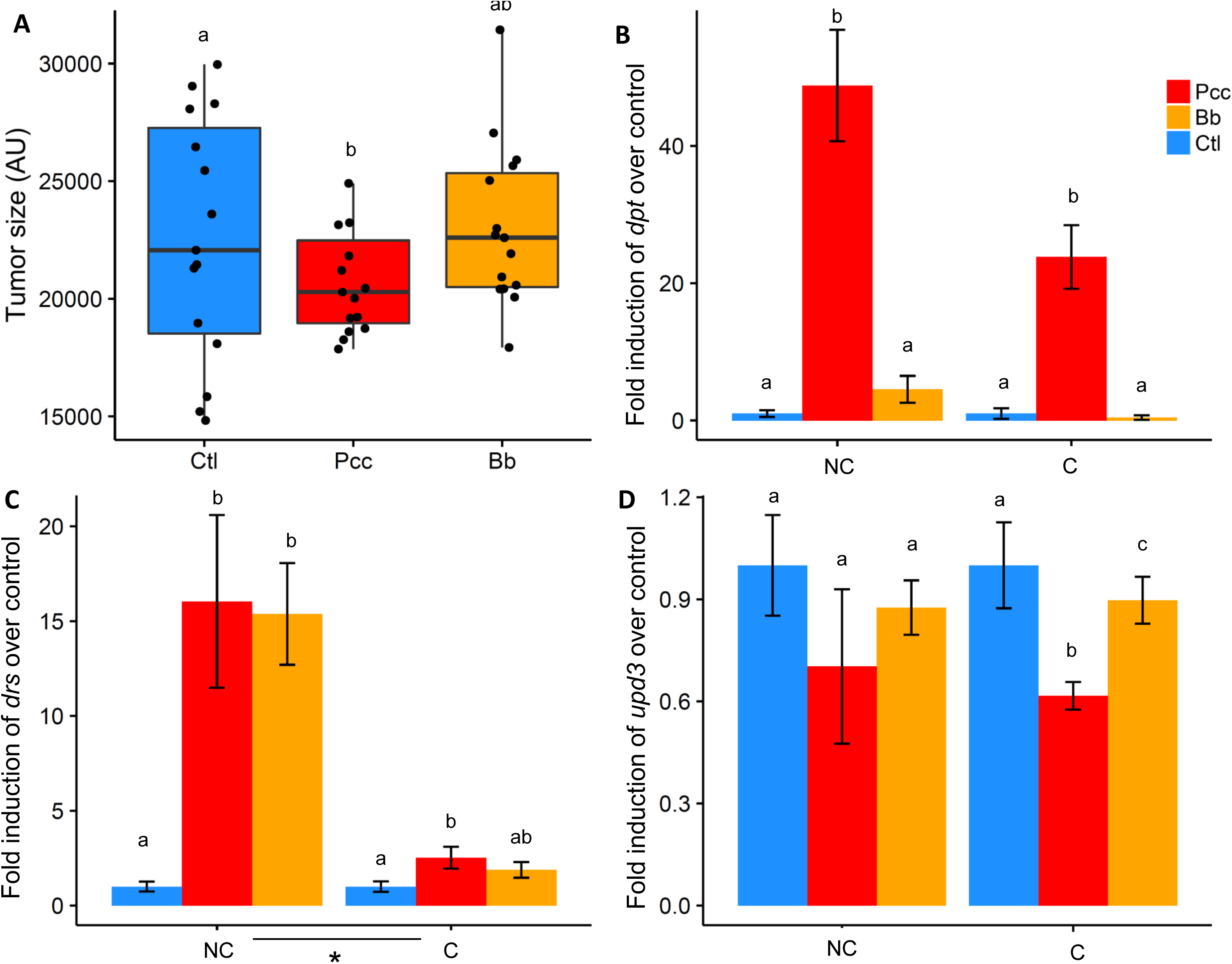
A) Effect of infection by bacterium *Pectobacterium carotovorum carotovorum* (abbreviated by *Pcc* in red) and the fungi *Beauvaria bassiana* (abbreviated *Bb* in orange) compared to control without infection (abbreviated Ctl in blue) on individual tumor size (S2 threshold: 75% of maximum intensity value). Fold induction of three immune genes: B) *dpt*, C) *drs* and D) *upd3* in three groups of larvae with different cancerous status (C: cancerous, G: non-cancerous) in response to infection by the bacterium *Pcc* (red) and the fungi *Bb* (orange) standardized by control without infection (blue). The error bars represent the standard error of mean relative expression calculated for all pools. Shared letters (a, b) above error bars denote means that are not significantly different, while different letters indicate that p<0.05 for pairwise comparisons (Tukey’s test). * represents a significant difference between status (p<0.05, status effect).

To identify which immune mechanisms could explain the effect of the bacterial infection, we compared the activation of the innate immune pathways following infections in cancerous versus non-cancerous larvae. Infection with *Pcc* resulted in at least a 20-fold increase of *dpt* expression compared to non-infected larvae of both statuses (Figure 2B; treatment effect: χ^2^ = 23.3, df = 2, pval < 0.0001). In non-cancerous larvae, *drs* expression was higher in *Pcc* and *Bb*-infected larvae compared to non-infected larvae (Figure 2C; treatment effect: χ^2^ = 12.9, df = 2, pval =0.002). However, the impact of the infectious treatment was comparatively lower in cancerous larvae (status effect: χ^2^ = 12.9, df = 2, pval=0.002).

Altogether, our results pointed out the role of Imd and Toll pathways in the tumor reduction observed in *Pcc*-infected larvae. Considering the recent studies demonstrating the anti-tumoral function of several antimicrobial peptides produced downstream of Imd and Toll pathway (*24, 25*), it is likely that tumor size reduction induced by bacterial infection was a consequence of a general activation of the innate immune system. However, Toll pathway alone seemed to be insufficient to induce a potent elimination of cancer cells, as suggested by the absence of tumor regression in *Bb*-infected larvae. This might be explained by the synergistic action of both pathways already reported for several antimicrobial peptides (*16*). Nevertheless, we cannot preclude that other mechanisms, such as metabolic or microbiota alterations consequent to digestive infection, were involved in tumor size reduction. In addition, the innate immune mechanisms, observed in response to *Pcc* infection, are susceptible to apply to other bacteria, including those encountered by humans, as they share common antigens called pathogen-associated molecular patterns (PAMPs) (*26*).

Finally, the expression of *upd3* gene was significantly decreased by *Pcc* and *Bb* infection compared to non-infected larvae but only in cancerous larvae (Figure 2D; z-value = −2.8, pval = 0.01). Interestingly, Jak/STAT pathway has been shown to have a dual role in tumoral immune responses. On the one hand, it promotes hemocytes proliferation, which subsequently restrict tumor growth (*19*) but on the other hand expression of *upd3* in the tumor has been shown to increase cancer cells proliferation (*23*). Thus, the downregulation Jak/STAT following infection could be directly involved in tumor size reduction or a by-product of cancer cell elimination.

Thanks to our experimental model we have been able to study the effects of one infectious event on cancer development. However, reports from Coley’s experiments as well as the use of bacillus Calmette-Guerin to treat superficial bladder cancer pointed out the importance of recurrent infections to promote efficient tumor regression (*6, 27*). Since it is experimentally challenging to do repeated infection on a short-lived model like *Drosophila*, we investigated the consequences of these different patterns of infection through a mathematical model that allows the exploration of a large number of infectious situations and where parameters have been estimated to reproduce the level of immune gene activation and tumor size observed experimentally (Supplementary materials). When this model simulated short repeated bacterial infections, we forecasted that it may result in a smaller tumor size compared to a single persistent infection (Fig. 3A). The effect of repeated infections can be linked to the maintenance of a higher level of Jak/STAT activation (Fig. 3B), which may act together with Imd and Toll pathways to eliminate cancer cells.

**Figure 3:**
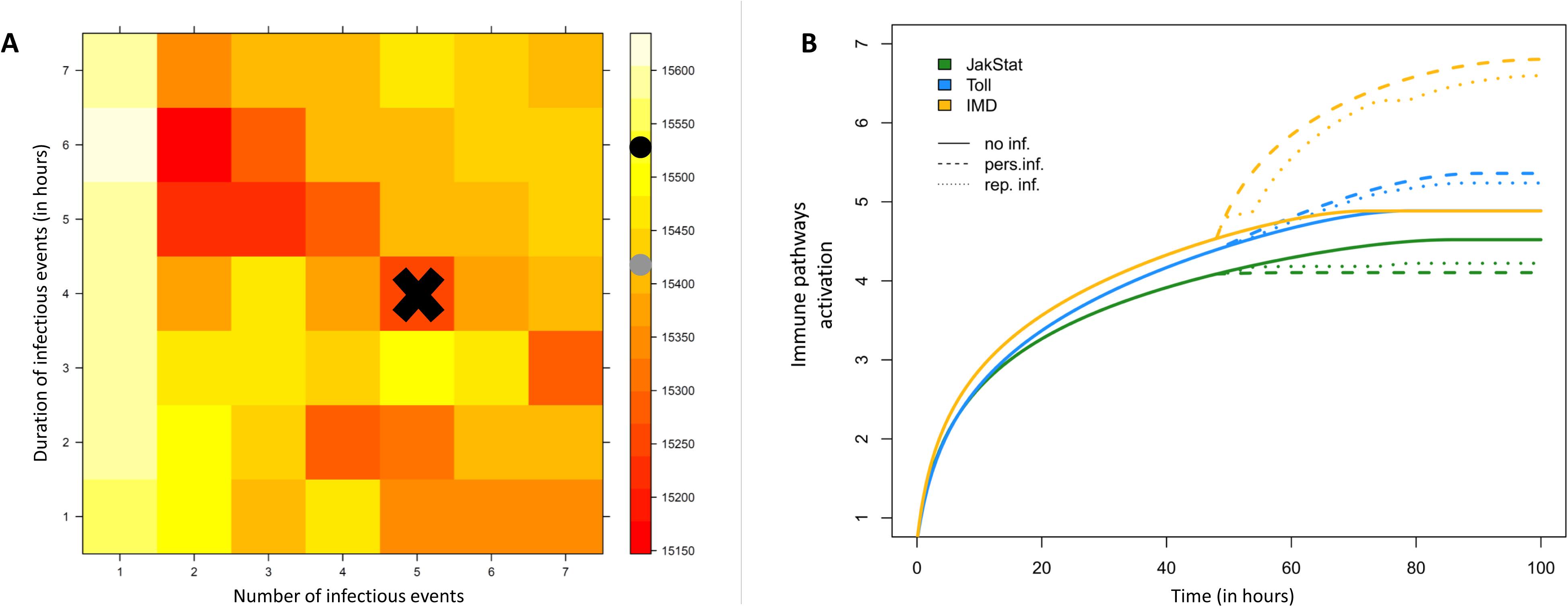
A) Influence of the number of infectious events and their duration on the accumulation of cancer cells (range from dark red for accumulation of <15200 cancer cells to white >15600 cells). Dark and gray dots represent the value obtained without infection and with a persistent infection respectively. The black sign shows the parameters used to obtained figure 4B. B) Mathematical model simulations exploring the activation of immune pathways (green represents Jak-STAT, blue Toll and yellow Imd) across times of experiments (in hours) in cancerous larvae without infection (in bold line), after repeated bacterial (dotted lines) or after persistent infection (dashed line). The parameters used to model dynamics of cancer cells and immune activation are in the supplementary materials (Table M3).

Our study demonstrates that naturally-occurring bacterial infection can promote tumor regression in *Drosophila* and that infection-dependent activation of the main innate immune pathways is likely to be a key driver of this regression. Since the 20^th^ century, radiotherapy and chemotherapy have been intensively developed to the detriment of the early attempts of immunotherapy, such as Coley’s toxin (*7*). However, the striking raise of immunotherapeutic strategies in the past decades, symbolized by the attribution of the 2018 Nobel Prize of Medicine, calls for a re-evaluation of the mechanisms linking infections and spontaneous tumor regression. While studies have mainly focused on adaptive immunity for the elaboration of new drugs (*28*), we focused on the role of innate responses. Our study, together with other recent studies in flies (*24, 25*), has revealed some molecular aspects mediating the forgotten potential of the innate immune system in the control of tumor burden. Further efforts should be made to elucidate the precise molecular mechanisms leading to cancer regression in response to innate immune system activation as it could open new perspectives to design treatments that mimics the effects of natural infections.

## Acknowledgments

We thank Jacques Montagne for providing us drosophila stocks. We thank Anna Dostolova for providing us Bb stocks. CJ would like to thank Laura Arenas and Marie-Lou Rollin for their help in infecting experiments. CJ thanks Virginie Georget from MRI plateform and Philippe Clair from qPHD plateform. This work was supported by the ANR (Blanc project EVOCAN to FT), by the CNRS (INEE), by André HOFFMANN (Fondation MAVA). All data and code to understand and address the conclusions of this research are available in the main text and supplementary materials.

## Supplementary Materials

Materials and Methods

Figures S1-S4

Tables S1-S2

