## Supplementary materials for "Infection triggers tumor regression through activation of innate immunity in *Drosophila*"

**This PDF file includes:**

Materials and Methods

Supplementary Text

Figs. S1 to S4

Tables S1 to S2

Materials and Methods

**Tumor model and drosophila stocks**

The genetically modified *Drosophila melanogaster* used in this study were engineered to develop a tumor of the eye disc, as previously described elsewhere (*1*). Briefly, the genetic scheme uses a truncated *eyeless* promoter-driven FLP recombinase expression (*ey(3.5)-FLP*) to generate discrete patches of GFP-labeled mutant cells specifically in the developing larval eye-antennal imaginal discs. Clones are mutant for the cell polarity regulator *scribble* (*scrib*) and mis-express a constitutive active form of ras oncogene (*Ras^v12^*). Male *yw;* Sp/Cyo; *FRT82B/TM6* flies were crossed with *yw, ey(3.5)-FLP; act5>stop>gal4,UAS-GFP; FRT82B, Tub-gal80* females to generate non-cancerous larvae (referred as NC) bearing GFP-labeled healthy clones. Male *UAS-Ras85^v12^; FRT82B, scrib^1^ /TM6* were crossed with the same females described above to generate cancerous larvae (referred as C) bearing *Ras^v12^/scrib^1^* GFP-labeled tumors. All Drosophila stocks and crosses were maintained on standard fly medium, under 12h/12h light dark cycles.

**Infectious agents**

*Pectobacterium carotovora carotovora* *Ecc15* (*Pcc*) is a gram negative bacterium that can naturally infect *Drosophila* (*2*). *Pcc* used in our study has been provided by the Pasteur Institut (Reference CIP 82.83T) and has been maintained by a shaking culture in LB medium at 29°C. *Beauvaria bassiana* (*Bb*) is a naturally occurring insect fungus, which infects flies during sporulation. *Bb* has been kindly provided by Anna Dostalova (École Polytechnique Fédérale de Lausanne, Switzerland). The fungus is grown on malt agar plates at 29°C in the dark for 2 weeks.

**Infecting experiments**

The crosses were allowed to lay eggs on sugar-agar plates for less than 24h, and collected embryos were incubated for two days at 24°C. From the two crosses, we selected larvae at the late 2^nd^ or early 3^rd^ instar stage based on the lack of the dominant Tubby phenotype carried on the *TM6* balancer chromosome.

An overnight culture of *Pcc* was pelleted by centrifugation (15 min at 4500 rpm) and adjusted to OD_600_ = 200. Oral infection was conducted 4 hours at 29°C by incubating larvae with 1.5mL of bacterial suspension mixed with crushed banana in small petri dishes according to the protocol described elsewhere (*2*). To realize cutaneous infection with *Bb*, larvae were rolled on spores and incubated in spores for 2h at 29°C (*3*, *4*). Then, larvae were transferred in crushed banana to avoid a bias caused by the diet change and incubated 2h more at 29°C. Finally, a control group was incubated in crushed banana without infectious agent for 4h at 29°C. After incubation, seven larvae were transferred into each well of a 96-well plate filled with yeast-sugar-agar medium and wells were plugged. The persistence of the infection was assessed in random groups of larvae (without selection for cancerous status) through classical PCR with specific primers (Table M1).

**Table M1**| Prevalence of infection at day one after exposure and persistence of infection two days after exposure.

|  | Prevalence | |
| --- | --- | --- |
|  | Day 1 | Day 2 |
| *Pcc*-infected larvae | 93.3% | 37% |
| *Bb*-infected larvae | 46.7% | 56.7% |

**Quantification of the tumor size**

After two days of incubation at 24°C post-infection, we performed tumor visualization of late 3^rd^ instar larvae using a dissecting microscope under GFP fluorescence (Zeiss A Lovert 200M, 2.5X).

Tumors were isolated with the software ImageJ 1.41.0 (*5*). Then, intensity values of each pixel have been standardized by the exposure time (see supplementary materials for more details). Finally, tumor size was quantified with Matlab (MATLAB 2015b, The MathWorks, Natick, 2015) as the number of pixels with scaled intensity above a given threshold. Four intensity thresholds were considered to measure tumor size: pixel intensity superior to 50% (S1), 75% (S2), 90% (S3) or 95% (S4) of the maximal intensity recorded in the picture (Figure M1). Thus, we obtained four distinct measures of the tumor standardized by exposure time that have been analyzed separately to quantify the robustness of our results.

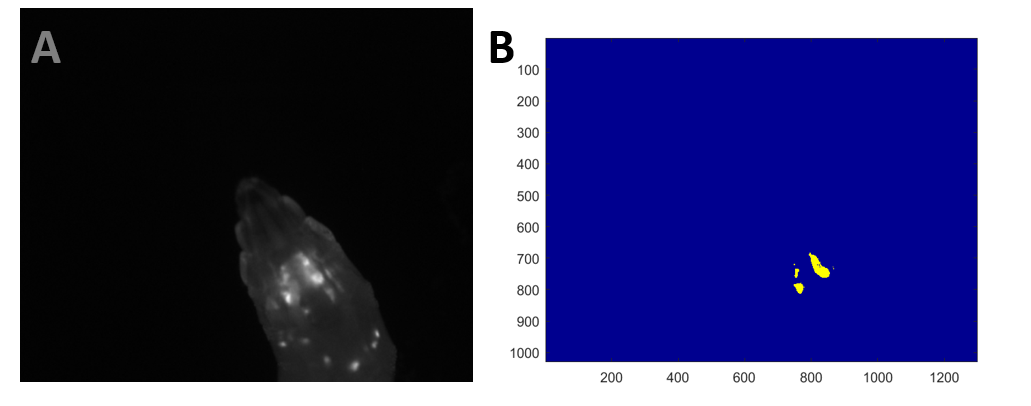

**Figure M1 | Visualization of GFP-labeled tumor in vivo**. A. A cancerous larva where we can observe GFP-labeled cells in eye-antennal discs. B. Number of pixels obtained after isolation and standardization with S1 threshold (i.e., pixel with an intensity superior to 50% of maximal intensity).

**qRT-PCR analysis**

After infections and determination of cancerous status, we obtained 10 pools of 7 larvae for each group of infection (non-infected, *Pcc* or *Bb* -infected larvae) for non-cancerous larvae. We obtained 15 pools of 7 larvae for each infectious treatment for cancerous larvae. All pools were fixed by liquid nitrogen and stored at -80°C. Total RNA was extracted using a TRIzol reagent following manufacturer’s protocol. Total RNA was eluted in 50µL of water and 25µL were used to perform DNase treatment (Turbo DNA-free kit, Life Technologies). Finally, total RNA was quantified using Nanodrop (ThermoScientific) and cDNA was synthetized from 2.5 µg of total RNA using SuperScript III kit (SS III First-Strand Synthesis System, Life Technologies) in 20µL total volume according to the manufacturer’s instructions.

We quantify expression levels of the *drosomycin* (*drs*), *diptericin* (*dpt*) and *unpaired 3* (*upd3*) genes using gene-specific primers (see Table M2). PCR reactions were conducted in a total volume of 3µL with LightCycle ® 480 SYBR Green I Master (Roche). 10 min pre-incubation at 95°C was followed by 40 cycles of amplification: 10s at 95°C, 10s at 64°C and 10s at 72°C. Melting curves were generated after the final amplification cycle by denaturating the amplicons at 95°C for 5s, cooling to 65°C for 1 min, and then increasing up to 97°C. Melting curves were used to estimate the specific melting temperature for each reaction. For each of the 5 genes, standard curve was generated from 10-fold serial dilutions of PCR amplicons. Each sample (*i.e*., pool) was analyzed in triplicates. The PCR efficiency and threshold cycle (Ct) were calculated by LightCycler ® 480 software. Only triplicates with a Ct standard deviation inferior to 0.5 were kept for the following analyses. Threshold cycle values were expressed relatively to two housekeeping genes (*rp49* and *tub84b*) (*6*) according to the procedure described in (*7*) which is based on the common method of 2^-ΔΔCt^. After normalization and inter-runs calibration, we obtain calibrated normalized relative quantity (henceforth called NRQ). We also measured the coefficient of variations (CV) and gene stability (M) for reference genes. We found that *rp49* and *tub84b* had a CV of 9.3 % and 8.9 % respectively which are considered as acceptable values (*i.e*., <25% (*7*)). M was equal to 0.26, which means that our reference genes were stably expressed and our samples homogenous.

**Table M2**| Primers used in this study

| **Gene names** | **Symbol** | **Forward (**5’-3’) | **Reverse (**5’-3’) |
| --- | --- | --- | --- |
| *diptericin* | *dpt* | GCTGCGCAATCGCTTCTACT | TGGTGGAGTGGGCTTCATG |
| *drosomycin* | *drs* | CGTGAGAACCTTTTCCAATATGATG | TCCCAGGACCACCAGCAT |
| *unpaired 3* | *upd3* | GCGGGGAGGATGTACC | GTCTTCATGGAATGAGCC |
| *ribosomal protein 49* | *rp49* | GACGCTTCAAGGGACAGTATCTG | AAACGCGGTTCTGCATGAG |
| *alpha-tubulin 84b* | *tub84b* | TGTCGCGTGTGAAACACTTC | AGCAGGCGTTTCCAATCTG |

**Statistical analyses**

We realized pairwise comparison through F-tests, T-test, or “Tukey” tests (using the “glht” function in package *multcomp*). Then, we tested the effect of tumor size, cancerous status and infectious treatments on immune profile through generalized linear mixed models (GLMMs), as implemented in the package *lme4* (*8*). Backward model simplification has been made according to the Akaike information criterion (AIC) (*9*). For the most parsimonious model, variable selection was conducted through analysis of variance (ANOVA) (using the ‘Anova’ function in package *car*, with test specified as “type3”) that tests the effect of each variable after all other factors have been accounted for (*10*). In addition, we used a principal component analysis (PCA) on immune genes expression to identify the relative contribution of each pathway. For each of the dimensions, we then calculated standard coordinates for pools by dividing the principal coordinates (*i.e*. loadings) by the square root of the dimension's eigenvalue. This standardization allowed them to be considered as independent variables. We used the standardized dimensions loadings as fixed factors in a GLMM explaining pooled tumor size. We concentrated our analyses on tumor size at pool level however results for individual data are available in supplementary materials. All the previous analyses were conducted using R v3.1.2 statistical software (R Development Core Team).

**Mathematical modeling**

In order to extrapolate what could be the consequences of different bacterial infection setups (*i.e.* repeated infections compared to persistent infection), we have developed a mathematical model aiming to reproduce our experimental results:

$$\frac{dC}{dt}=rC\left[ 1-\frac{C}{K} \right]-{C(\alpha}_{jc}J+\alpha_{tc}T+\alpha_{ic}I)$$

$$\frac{dJ}{dt}=R_{j}+ \beta_{j}\left[ 1-\frac{J}{\Phi} \right]+\varepsilon_{ij}I+\varepsilon_{tj}T+\theta_{cj}C$$

$$\frac{dI}{dt}=R_{i}+ \beta_{i}\left[ 1-\frac{I}{\Phi} \right]+\varepsilon_{ji}J+\varepsilon_{ti}T+\theta_{ci}C$$

$$\frac{dT}{dt}=R_{t}+ \beta_{t}\left[ 1-\frac{T}{\Phi} \right]+\varepsilon_{jt}J+\varepsilon_{it}I+\theta_{ct}C$$

In this model J, I and T represent the activation of Jak-STAT, IMD and Toll pathways respectively. Cancer cells (C) replicate at rate *r* with a maximal total number of cells *K* (*i.e*. carrying capacity) and they are eliminated by the three immune pathways at rate $\alpha_{*c}$ (_*_ represents the immune pathway considered). Each pathway has a basal activation level $R_{*}$. As we assumed that immunity is controlled to avoid auto-immunity, activation cannot overpass a maximal value$\Phi$. In a similar manner, we assumed that reciprocal interactions exist among immune pathways and are characterized by the matrix$\varepsilon$. Each pathway can be activated by cancer cells at rate $\theta_{c*}$ and by bacterial infection at rate *β*. All parameters have been chosen in accordance with the literature or to reproduce our experimental protocol (where infection occurs at half lifetime, *i.e*., 50 hours after egg laying until the end of the experiments, *i.e*., 100 hours) and our experimental results.

The first step of our modeling approach was to estimate the unknown parameters in order to create a parsimonious model able to reproduce the experimental data (Table M3). To this end, we minimize the least-squares through the Nelder-Mead algorithm between experimental data and model outcomes for the activation of immune pathways and tumor size 48H after infection beginning. We ran our model from infection time point to the end of the stimulation with initials values estimated from our experimental data (C=300, J=8.6, T=150, I = 374). With this model, we were able to reproduce our experimental results (Fig. M3). Even if it slightly underestimates the activation of Imd following bacterial infection, all the other variables have been successfully reproduced. Finally, we used this fitted model and estimated parameters to assess the effect of repeated infections (i.e., 5 infectious events of 4h in 48h) on tumor cell accumulation. We started all our simulations by considering that larvae have at least one cancer cell (because crossing of the two fly strains is sufficient to generate tumor cells) and harbor very few immune effectors (C=1; J=1; I=1; T=1).

**Table M3|** Parameters value used to model dynamics of cells and immune activation.

| **Parameter** | **Method** | **Description** | **Value used** | **Justification / Interpretation** | **References** |
| --- | --- | --- | --- | --- | --- |
| $R$ | Fixed | Basal activation | $R_{i}$= 0.46  $R_{j}$=0.215  $R_{t}$=0.13 | Calculated with the formula of logarithmic growth: N’(t)=rN(t)*(1-N(t)/K) |  |
| *K* | Fixed | Carrying capacity of the eye disc | 22000 | Calculated from observed data. Maximal value in non-infected larvae + 10% |  |
| $\Phi$ | Fixed | Maximal immune activation level | 2e^6^ | Calculated from observed data. Maximal value all pathways confounded + 10% |  |
| $\beta_{t}$ | Fixed | Activation of Toll by bacterial infection | 1.9 | Based on fold induction at 12h after infection compared to non-infected. | (*11*) |
| $\beta_{i}$ | Fixed | Activation of Imd by bacterial infection | 20 | Based on fold induction at 12h after infection compared to non-infected. | (*12*) |
| $\beta_{j}$ | Estimated | Activation of Jak/STAT by bacterial infection | -1.6e^-02^ | Bacterial infection could directly down-regulate Jak/STAT pathway. |  |
| $r$ | Estimated | Replication rate of cancer cells | 0.1 | By hours |  |
| *α_jc_* | Estimated | Elimination of cancer cells by Jak/STAT pathway | 1.8e^-04^ | Jak/STAT promotes hemocytes proliferation. | (*13*) |
| *α_ic_* | Estimated | Elimination of cancer cells by Imd pathway | 2.6e^-05^ | *Defensin* has been shown to eliminate cancer cells. | (*14*) |
| *α_tc_* | Estimated | Elimination of cancer cells by Toll pathway | -1.6e^-04^ | Toll pathway could have a pro-tumoral role. Ortholog of the pathway activate by NF-κB implied in cell proliferation and angiogenesis | (*15*) |
| θ*_tc_* | Estimated | Activation of Jak/STAT by cancer cells | -6.6e^-04^ | Cancer cells are known to generate an immunosuppression. | (*16*) |
| θ*_jc_* | Estimated | Activation of Toll by cancer cells | -1.6e^-04^ |  |  |
| θ*_ic_* | Estimated | Activation of Imd by cancer cells | -1.3e^-03^ |  |  |
| *ε_ti_* | Estimated | Impact of Toll on Imd | 2.8e^-03^ | Imd can influence the Toll pathway through the control of PGRP-SA. | (*17*) |
| *ε_it_* | Estimated | Impact of Imd on Toll | 4.3e^-03^ |  |  |
| *ε_tj_* | Estimated | Impact of Toll on Jak/STAT | 4.5e^-05^ | Toll can control the JAK/STAT cascade. | (*18*) |
| *ε_jt_* | Estimated | Impact of Jak/STAT on Toll | 1.2e^-02^ |  |  |
| *ε_ji_* | Estimated | Impact of Jak-STAT on Imd | 1.3e^-02^ | JNK (activated during Imd cascade) inhibition resulted in a reduction of JAK/STAT reporter levels | (*19*) |
| *ε_ij_* | Estimated | Impact of Imd on Jak/STAT | -2.8e^-04^ |  |  |

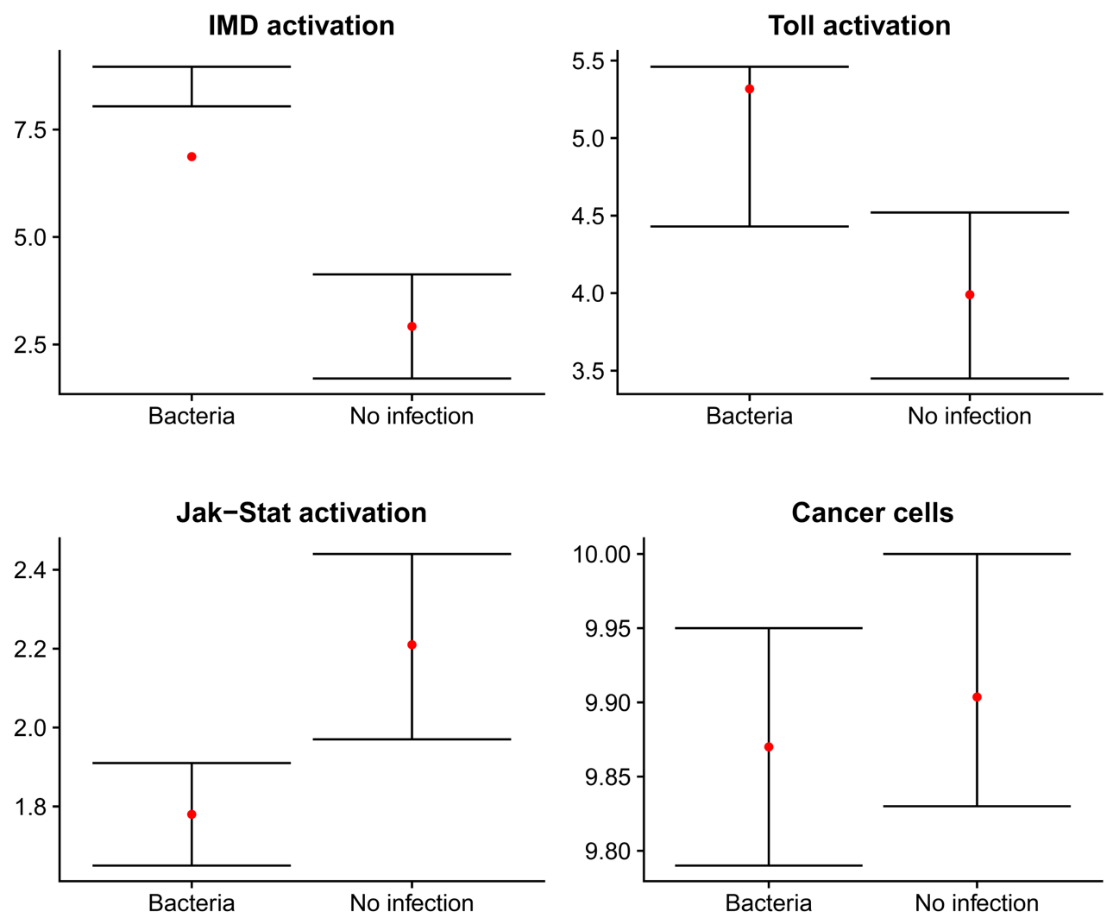

**Figure M2|** Reproduction of observed experimental data with simulated data obtained with an ordinary differential equation model using parameters estimated by optimization method. Confidence interval of observed data are in black and red dots represent simulated data. Value in the y-axis have been log-transformed.

**Supplementary Text**

**Tumor visualization**

Before correction, pixel intensities were comprised between 1 and 4000 (Fig. S1). Regarding image analyses, we noticed that the intensity of halogen lamp was not correlated with tumor size (see Fig. S2A) and thus we did not consider this factor in our analyses. However, we found that exposure time significantly impact tumor size estimation and thus we standardized our tumor measurements by the exposure time. This correction allowed an absence of correlation between tumor size and exposure time (Fig. S2B).

In order to control the effect of infectious treatments on tumor size through a modification of larval size, we also measured the mouth hook as a proxy of body size (Mirth et al. 2005) in ImageJ software. We looked for an effect of treatment on hook size with a specific linear model and realized pairwise comparisons of body size between groups using t-tests. Then, for each quantification of tumor size (S1 to S4), we have designed GLMMs where tumor size was used as the response variable, hook size as fixed factors and the exposure time as a random factor. We fitted each model to the most parsimonious distribution and conducted variable selection as described in the methods. The effect of body size was not significantly different depending on infectious treatment and tumor size was not affected by body size (see Fig. S3A and B).

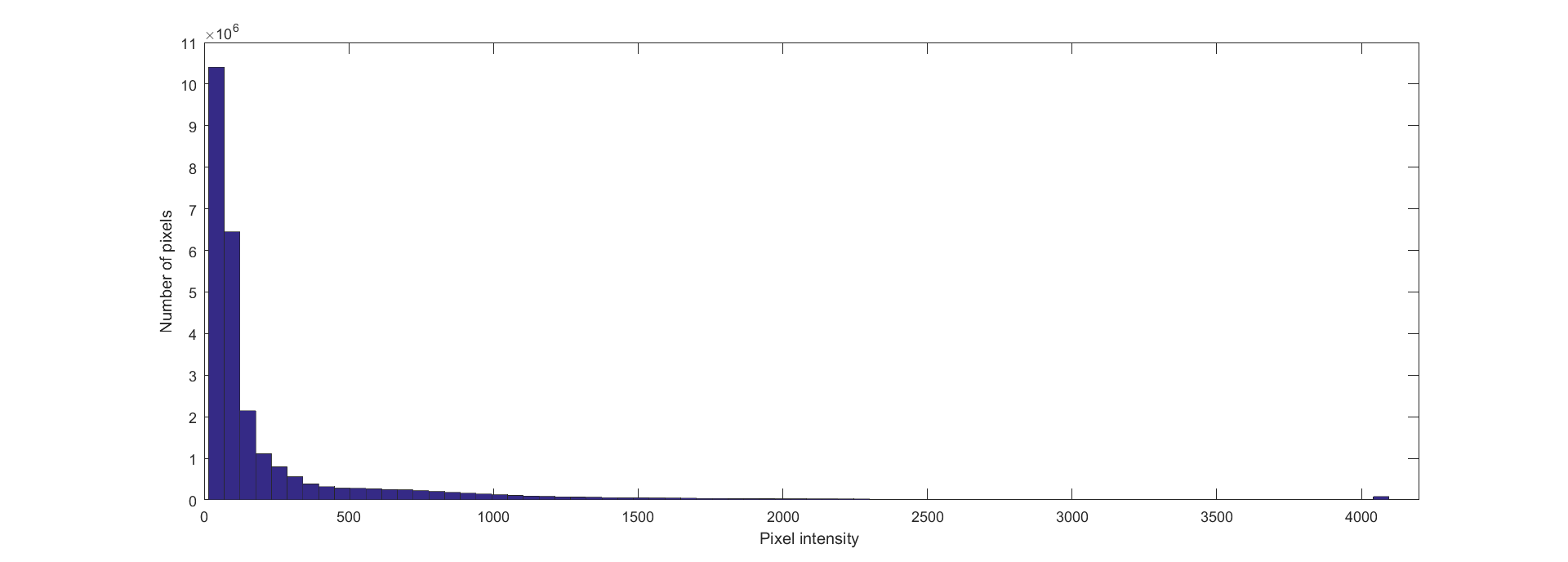

**Figure S1| Histogram of pixel intensities for each of the 759 tumor pictures**. The total number of pixels considered was above 25 billion and the figure represents data before transformation.

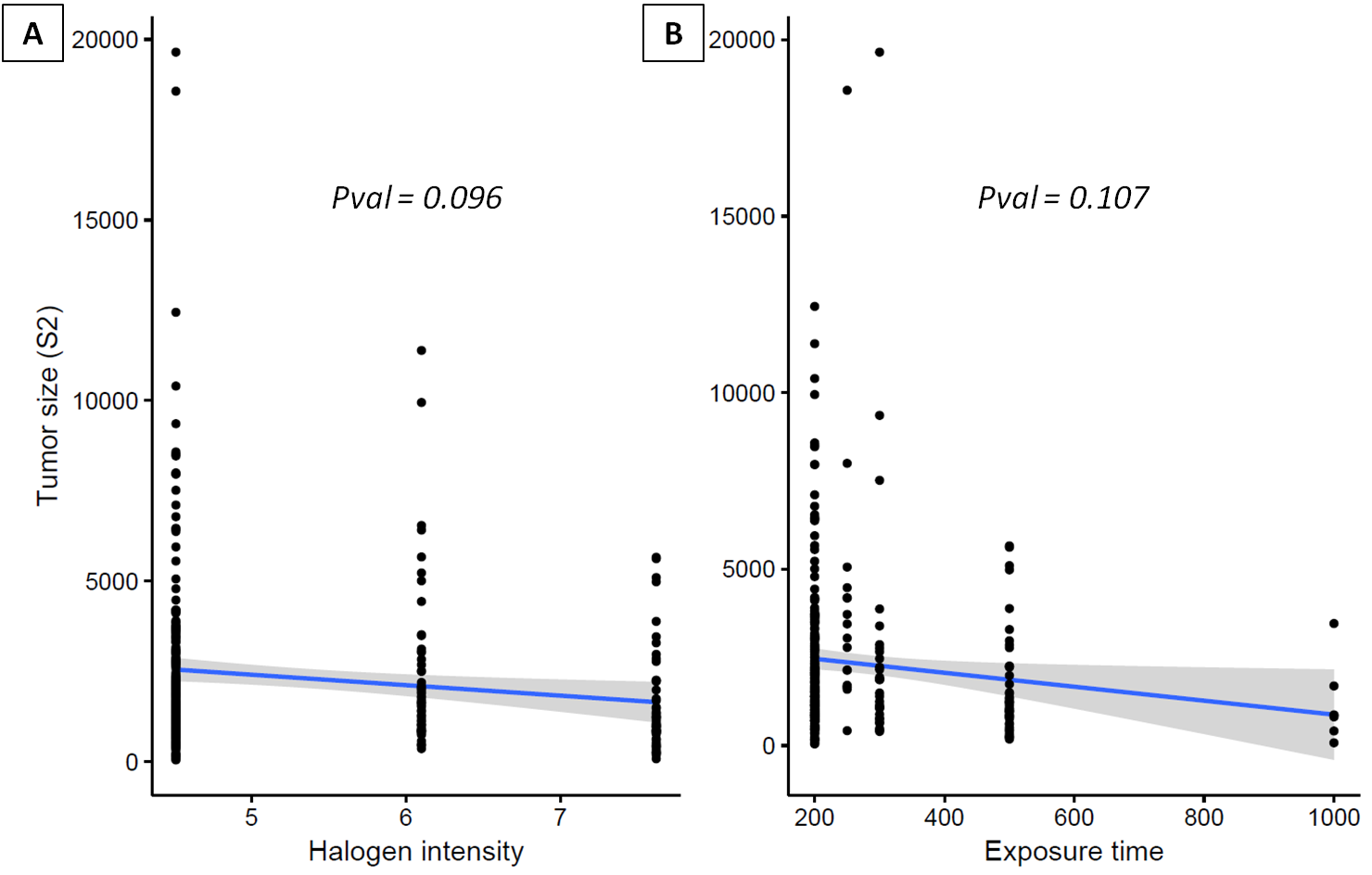

**Figure S2| Picture characteristics.** A) Absence of correlation between halogen intensity of GFP lamp and tumor size. B) Absence of correlation between exposure time and standardized tumor size.

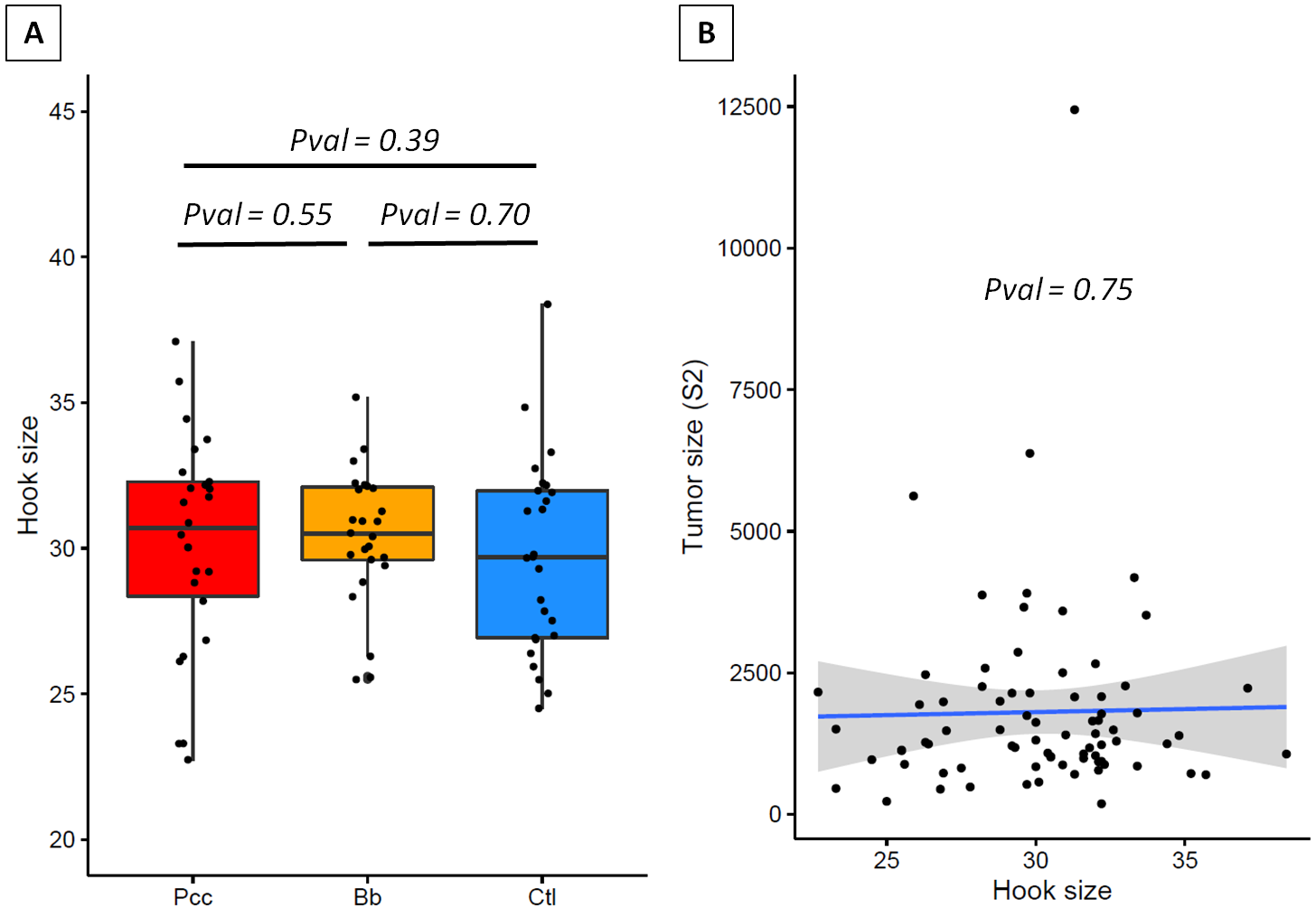

**Figure S3| Impact of body size on tumor size depending on infectious treatment**. A) Body size, approximate by mouth hook size, is not affected by infectious treatment. B) Tumor size is not associated with body size.

**Complementary analyses at individual level and with different tumor thresholds**

Information on tumor size and infectious treatment were available at individual level before larvae have been pooled for transcriptomic analysis. A GLMM on individual data was constructed with individual tumor size used as response variable, treatment as fixed factor and replicates as random factors. The principal results from tumor size analyses depending on the infectious treatment are presented in Table S1 and they are compared to those obtained at pool level.

**Table S1**| Results from tumor size analyses. W: Mann-Whitney statistic; T: t-test statistic; F: fisher test statistic.

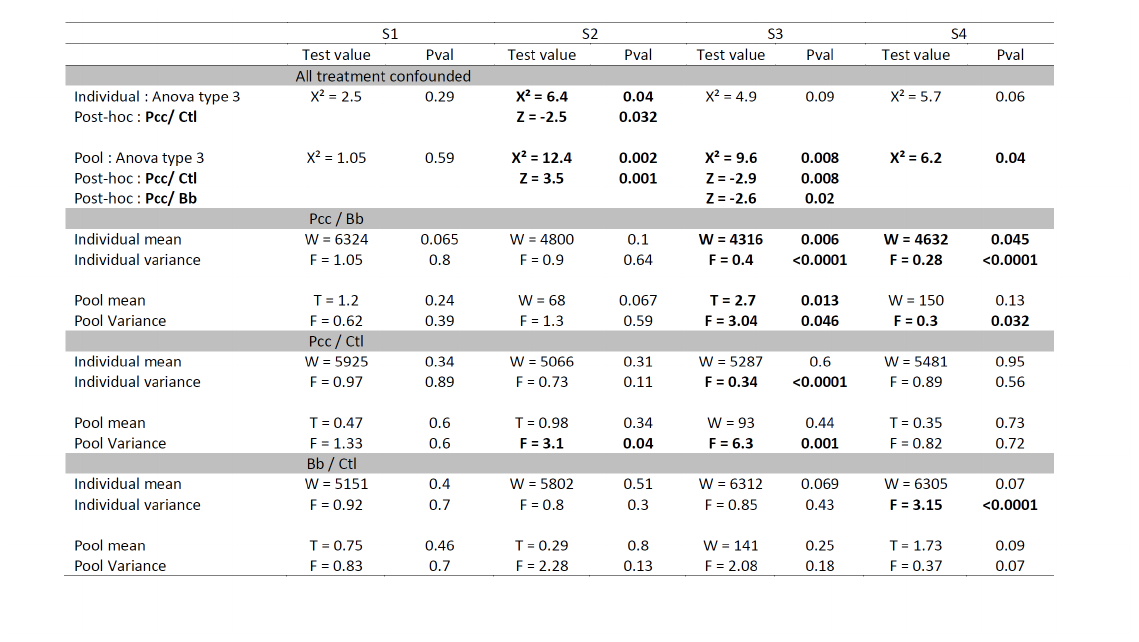

**Supplementary materials references**

10.	J. Fox, S. Weisberg, *An {R} Companion to Applied Regression, Second Edition.* (Thousand O., 2011).
